# Allosteric factors in the calcium/calmodulin-responsive kinase II hub domain determine selectivity of GHB ligands for CaMKIIα

**DOI:** 10.1101/2025.01.19.633782

**Authors:** Stine Juul Gauger, Line B. Palmelund, Yongsong Tian, Aleš Marek, Mathias R Namini, Nane Griem-Krey, Stefanie Kickinger, Jonas S. Mortensen, Bente Frølund, Petrine Wellendorph

## Abstract

The Ca^2+^/CaM-dependent protein kinase II alpha (CaMKIIα) is a highly important synaptic protein, which comprises a unique holoenzyme structure organized via the central hub domain. Recently, a distinct binding pocket in the CaMKIIα hub domain was identified for the endogenous neuromodulator γ-hydroxybutyric acid (GHB) and related synthetic analogues. Key interacting residues in CaMKIIα were revealed, but the pronounced selectivity towards the *alpha* variant of CaMKII has remained unresolved. Aimed at elucidating the molecular determinants for this selectivity, we here conducted binding studies to CaMKII-HEK whole-cell homogenates using two different in-house-developed GHB-related radioligands, [^3^H]HOCPCA and [^3^H]O-5-HDC, in combination with site-directed mutagenesis. Binding to CaMKIIα with the smaller-type radioligand [^3^H]HOCPCA validated key involvement of the four known residues (His395, Arg433, Arg453 and Arg469), but also revealed a role for the upper hub flexible loop containing the CaMKIIα-specific residue Trp403 (Leu in all other CaMKII isozymes). Insertion of the corresponding residues (L467W/C533R) into CaMKIIβ failed to introduce [^3^H]HOCPCA binding. However, with the larger-type radioligand, [^3^H]O-5-HDC, specific binding in CaMKIIβ (L467W/C533R) was achieved. Thus, of the four native CaMKII isozymes, only CaMKIIα accommodates GHB ligands. The study identifies the CaMKIIα flexible pocket loop as a distantly located “allosteric” factor in determining selectivity of GHB analogues for CaMKIIα. It sheds light on a remarkable interplay of the entire hub cavity for accommodation of ligands, and corroborates GHB analogues as CaMKIIα-selective.

## Introduction

The Ca^2+^/CaM-dependent protein kinase II (CaMKII) is a Ser/Thr kinase involved in several Ca^2+^-signalling pathways. Four distinct, but highly related, genes (α, β, γ and δ) encode the four main isozymes of CaMKII^1^. CaMKIIα and -β are predominantly found in the brain and play an important role in synaptic plasticity underlying learning and memory (reviewed by^2,3^). CaMKIIδ and -γ are more ubiquitously expressed and are found in the brain, skeletal muscle and the heart, with CaMKIIδ suggested to be involved in cardiovascular diseases^4^. CaMKII has a unique holoenzyme structure composed of subunits assembled into a stacked arrangement of two donut-shaped structures^5,6^. Each subunit consists of an N-terminal catalytic kinase domain, a regulatory segment containing the binding site for Ca^2+^/CaM and central phosphorylation sites (Thr286 and Thr305/Thr306), a variable linker, and a C-terminal hub domain responsible for holoenzyme assembly^5^. The hub domains of CaMKIIα and -β share 77% overall sequence identity^6,7^. In addition to the canonical holoenzyme assembly, a role for the hub domain in allosteric regulation of kinase activity is emerging, supported by structural work^5,7,8^. Furthermore, the hub via its known fluctuations in oligomeric structure, is suggested to play a role in spreading of kinase activity (activation-triggered subunit exchange), or may occur as inter-holoenzyme phosphorylation without hub domain mixing^7,9–11^ potentially mechanistically involved in sustainment of memories^12,13^.

In 2021, we reported the identification of a molecularly distinct binding pocket in the hub domain of CaMKIIα, which specifically binds small molecules related to the γ-aminobutyric acid metabolite, γ-hydroxybutyric acid (GHB)^14^. This work was enabled by the availability of selective and nanomolar affinity GHB analogues^15–17^. Intriguingly, GHB analogues bind exclusively to CaMKIIα, corroborated by the complete absence of binding in brain slices from *Camk2a^-/-^* mice with three different GHB ^3^H-labelled ligands, and a photoaffinity ligand, yet binding was fully preserved in *Camk2b^-/-^* tissues^14^. Also, radioligand binding was absent in HEK-cell expressed CaMKIIβ, γ and δ^14^. Using the high-affinity analogue, 5-hydroxydiclofenac (5-HDC) (Fig. 1A)^17^, a crystal structure of the hub domain was obtained, highlighting the key residues involved in the binding pocket. These key residues included the positively charged Arg433, Arg453 and Arg469, and His395^14^. Additionally, the crystal structure revealed a significant movement of Trp403 (Trp-flip) upon binding of 5-HDC^14^. This outward placement of Trp403 has since been corroborated with other GHB ligands^18,19^. Interestingly another key tool compound, 3-hydroxycyclopent-1-enecarboxylic acid (HOCPCA) (Fig. 1A), does not cause a detectable Trp-flip, probably due to its small size^14^. Hence, the flexible loop in the upper cavity of the identified binding pocket (the pocket loop) can present with two distinct conformations: An inward-flipped conformation with the Trp403-associated loop pointing into the hub cavity, and an outward-flipped conformation where the Trp403-associated loop is displaced outwards. Intriguingly, GHB and brain-permeable analogues, such as HOCPCA, have been found to be neuroprotective in several mouse models of ischemia, pointing to a potential clinical utility in targeting the CaMKIIα hub domain^14,20,21^.

**Figure 1:**
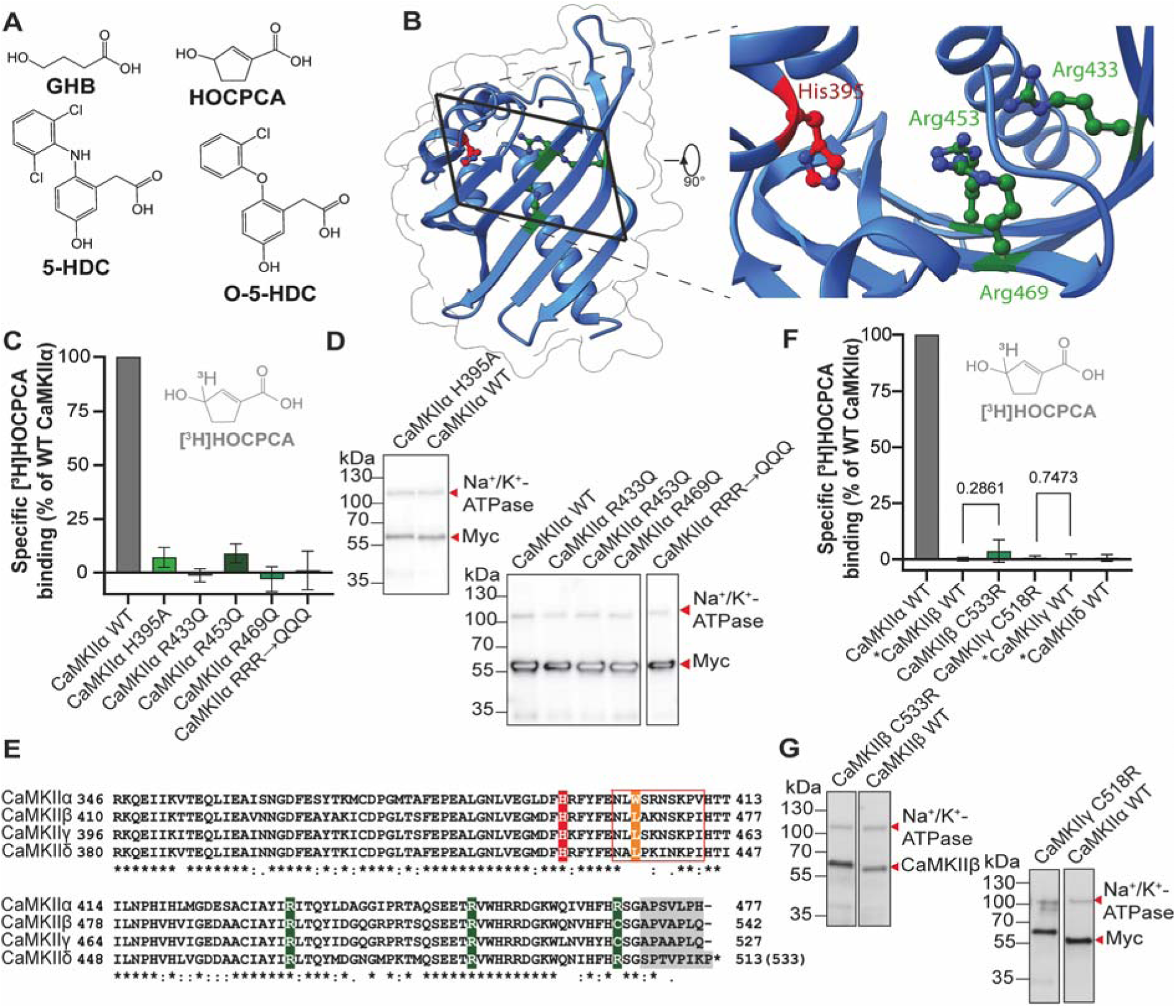
Molecular determinants for [^3^H]HOCPCA binding. **A)** Chemical structures of GHB ligands. **B)** Crystal structure of the CaMKIIα hub domain with the four core residues His395 in red, and Arg433, Arg453 and Arg469 in green. Close-up view shows the position and orientation of the residues towards the inside of the binding pocket (PDB: 7REC). **C)** Binding of [^3^H]HOCPCA to CaMKIIα binding pocket mutants; RRR→QQQ, triple mutant R433Q/R453Q/R469Q. **D, G)** Representative western blots confirming expression; Na^+^/K^+^-ATPase as loading control. **E)** Sequence alignment of CaMKII rat hub domains with key molecular determinants highlighted. The red box signifies the pocket loop with the key residue W403 in orange; predicted C-tail in grey. **F)** Specific binding of [^3^H]HOCPCA to CaMKIIβ WT, CaMKIIβ C533R, CaMKIIγ WT, CaMKIIγ C518R and CaMKIIδ WT. *Data taken from^14^. Data were pooled from three independent experiments performed in technical triplicates and shown as mean bar graphs ± SD. Student’s *t* test; significance level P<0.05.

So far, the pronounced selectivity of these small-molecule GHB ligands for CaMKIIα over CaMKIIβ, -γ or -δ has remained a riddle. This is especially intriguing as the majority of key residues identified in the binding pocket are either fully (CaMKIIδ) or partially conserved (CaMKIIβ and γ). Thus, this study sought to identify molecular determinants responsible for the CaMKIIα selectivity of GHB analogues by exploring specific residues both in the known binding pocket and the nearby cavity. To this end, we performed molecular modelling and site-directed mutagenesis of CaMKII isozymes. Engineered mutants were heterologously expressed in HEK293T cells and evaluated in radioligand binding assays with the standard radioligand [^3^H]HOCPCA, and a newly developed probe [^3^H]O-5-HDC based on chemical optimization of 5-HDC^17^.

## Materials and methods

### Compounds and radioligands

GHB sodium salt was purchased from Sigma-Aldrich (St. Louis, MO, USA). HOCPCA (3-hydroxycyclopent-1-enecarboxylic acid) sodium salt and 5-HDC (5-hydroxydiclofenac) were prepared as reported^14,17^ (Fig. 1A). [^3^H]HOCPCA (spec. activity 28.6 Ci/mmol) was produced in-house as described^22^. [^3^H]O-5-HDC (spec. activity 48.2 Ci/mmol) was produced as described in Supporting Information.

### Plasmids and mutants

For transfections we employed the following plasmids (all from Origene, MA, USA): pCMV6-CaMKIIa-Myc-DDK, (#RR201121), pCMV6-CaMKIIg-Myc-DDK (#RR207416), and pCMV6-CaMKIId-Myc-DDK. The pCAGG-CaMKIIb-pPGK-tdTOMATO plasmid for CaMKIIβ was a kind gift from Dr. G. van Woerden, and has been described previously^23^. Site-directed mutagenesis and verification of the sequences (rat) was performed by Genscript (Piscataway, NJ, USA). Intact expression of mutants was validated by western blot analysis as detailed in Supporting Information.

### Radioligand binding assays

Preparation of whole-cell homogenates including cell-culturing details, and preparation of cortical membrane homogenates are detailed in the Supporting Information.

Recombinant HEK293T whole-cell homogenate equilibrium binding experiments were performed similar to that described previously^14^, using a 48-well assay format. Briefly, whole-cell homogenate (100-150 µg protein per well) and either 40 nM [^3^H]HOCPCA or 10 nM [^3^H]O-5-HDC were incubated in a total volume of 400 µL binding buffer (50 mM KH_2_PO_4_, pH 6.0) for 1 hr on ice. Non-specific binding (NSB) was determined using 16-30 mM GHB. The binding reaction was terminated by precipitation with ice-cold acetone (1 hr at -20 °C), and radioligand-bound protein separated from free radioligand by vacuum-filtration through glass fibre (GF/C) filters. Counts (DPM) were detected by liquid scintillation counting^14^. Counts for CaMKIIα WT typically ranged from 2,000-6,000 DPM with [^3^H]HOCPCA and 1,800-5,000 DPM with [^3^H]O-5-HDC, slightly lower for the CaMKIIβ C657R/L790W mutant. All experiments were performed in technical triplicates using at least three different batches of HEK293T cell homogenates.

Characterization of [^3^H]O-5-HDC binding to native CaMKIIα (rat cortical homogenate) was carried out using a 96-well format assay based on a protocol very similar to [^3^H]HOCPCA^22^, see Supporting Information.

*In vitro* autoradiography was performed on 12-μm thick coronal mouse brain sections from adult male *Camk2a* (*Camk2^atm3Sva^*, MGI:2389262) (-/-) or corresponding litter mates (+/+) backcrossed in the C57BL/6j background. Experiments were performed as previously described^14^ using 0.09 nM [^3^H]O-5-HDC.

### Data analysis and statistics

Data analysis was performed on pooled, normalised data using GraphPad Prism 8 (GraphPad Software Inc., San Diego, CA, USA) as further detailed in Supporting Information. For statistical analysis, a Student’s *t* test or One-way ANOVA followed by Dunnett’ multiple comparison test for comparison of mean between groups was used, as specified in the figure legends. In a few instances a one sample *t* test with a hypothetical mean value of 100 representing normalised specific binding of WT CaMKIIα was used.

### Computational modelling

Protein visualization and alignment was performed in ChimeraX 1.7.1. The protein structures were loaded and single subunits from each PDB file were sequentially aligned using the MatchMaker extension of ChimeraX^25^. The pairwise sequence alignment, constructed by MatchMaker to guide the protein superposition utilizes the Needleman-Wunsch algorithm for global and local alignment. The superposition of the listed CaMKIIα hub domains was aligned with RMSD scores ranging from 0.969 Å to 1.461 Å. Multiple sequence alignment was performed with Clustal Omega (EMBL-EBI).

## Results and discussion

### Core molecular determinants for the binding of GHB ligands to CaMKIIα

The 5-HDC/CaMKIIα hub crystal structure (PDB: 7REC) revealed four key residues interacting directly with the ligand, thus constituting the core binding pocket: His395, Arg433, Arg453 and Arg469^14^ (Fig. 1B). These *core* molecular determinants are oriented with the sidechains directed towards an inner cavity of the binding pocket (Fig. 1B), enabling electrostatic and hydrogen bond interactions with 5-HDC. Thus, using site-directed mutagenesis of these four residues, we initially evaluated [^3^H]HOCPCA binding in CaMKII-HEK whole-cell homogenates. Convincingly, mutation of Arg433 or Arg469 into their non-charged counterparts (R433Q and R469Q), completely abolished [^3^H]HOCPCA binding (Fig. 1C). For the H395A and R453Q mutants, a very minor degree of specific binding remained (7-9%). As expected, the triple mutant (RRR433/453/469QQQ) was completely binding-deficient. Western blot analysis (Fig. 1D) confirmed intact expression of all mutants. Collectively, these data validate that the four residues, originally identified in the crystal structure of 5-HDC, are also essential for [^3^H]HOCPCA binding, thus resembling the key core molecular determinants for binding of GHB analogues to the CaMKIIα hub.

### Introduction of the missing arginine in CaMKIIβ and -γ fails to introduce GHB ligand binding

We previously showed that [^3^H]HOCPCA and related GHB analogues display remarkable selectively for CaMKIIα^14^. A simple explanation for the pronounced *alpha* selectivity could be the absence of one or more of the identified core molecular determinants. According to a multiple sequence alignment of the four CaMKII hub domains (Fig. 1E), among the four core determinants (marked green and red in Fig. 1E), only Arg469 (CaMKIIα) differs between the CaMKII isozymes. All four residues are conserved to CaMKIIδ, whereas Arg469 is a cysteine in CaMKIIβ and -γ (C533 and C518, respectively). Reasoning that introduction of the “missing arginine” in CaMKIIβ and -γ might restore [^3^H]HOCPCA binding, we introduced an arginine into the corresponding position in CaMKIIβ (C533R) and CaMKIIγ (C518R), and performed radioligand binding with [^3^H]HOCPCA. As shown in figure 1F, no specific binding was observed in either construct (Western blot analysis verified similar protein expression for all constructs (Fig. 1G)). This highlights that the mere presence of all four core binding residues is insufficient to introduce GHB ligand binding, and suggests additional molecular determinants to be responsible for the unique *alpha* selectivity.

### Importance of the CaMKIIα pocket loop for GHB ligand binding and selectivity to CaMKIIα

Next, we broadened the selectivity analysis by including regions and/or residues that might be important for binding in the nearby hub cavity. For this purpose, we aligned five available crystal structures of CaMKIIα hub proteins reported to have the lowest number of unresolved residues (Table S1). The overlay revealed two highly flexible regions in the hub domain with a fairly high degree of non-conserved residues, that were located opposite to each other at the upper part of the binding cavity (Fig. 2A). We hypothesised that one or both of these regions might play an important role for ligand accessibility and create a favourable environment for the bound ligand. In this study, we will refer to these two regions as the pocket loop and the C-tail. In CaMKIIα, the pocket loop consists of residues between the N-terminal alpha helix and the first beta sheet (residues from Asn401 to Val410), highlighted in the sequence alignment (Fig. 1E).

**Figure 2:**
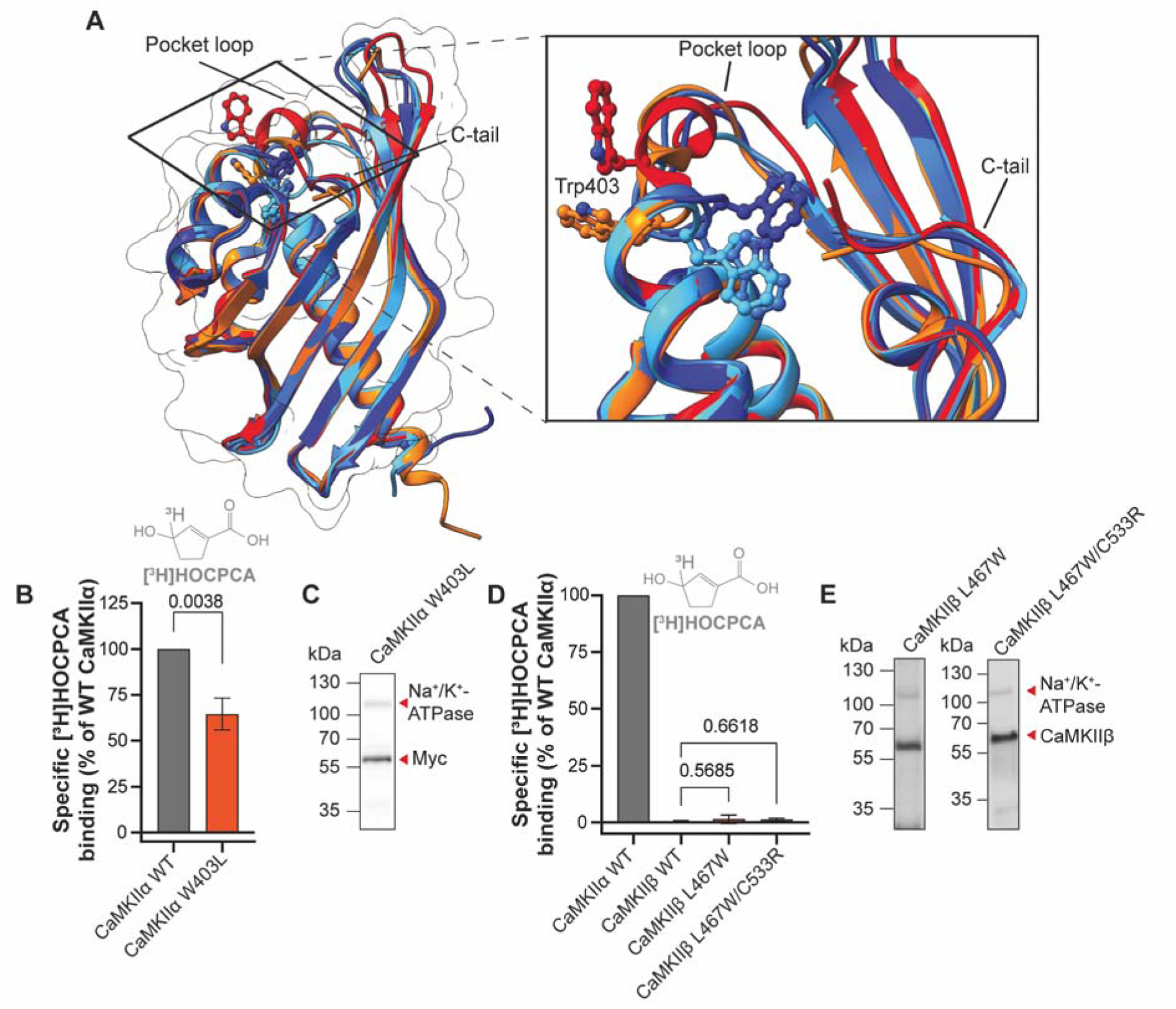
The CaMKIIα pocket loop is important for optimal binding of [^3^H]HOCPCA. **A)** Crystal structure alignment of CaMKIIα hub domains; unbound state (PDB codes: 7REC in blue, 3SOA in dark blue and 6OF8 in light blue), and bound state (PDB codes: 1HKX in orange and 7REC in red), highlighting the pocket loop and the C-tail. The close-up view shows the different orientations and positions of Trp403 in the loop. **B)** Specific [^3^H]HOCPCA binding to CaMKIIα W403L. One sample *t* test. **C, E).** Western blot validation of protein expression; Na^+^/K^+^-ATPase as loading control. **D)** Specific [^3^H]HOCPCA binding to CaMKIIβ WT, L490W and L490W/C533R. One Way ANOVA, post hoc Dunnett’s test. Significance level P<0.05. Data are pooled from three independent experiments performed in technical triplicates and shown as mean bar graph ± SD.

In the structural overlay (Fig. 2A), large conformational flexibility was observed for Trp403 in the pocket loop, a residue unique to CaMKIIα (Fig. 1E). As demonstrated, this Trp403 moves substantially from the inside of the cavity to the outside upon occupancy of the pocket (Fig. 2A)^14,18,19^, and appears to depend on the size of the ligand and its ability to extend into the upper part of the pocket^19^. Interestingly, in one reported structure (PDB 1HKX), binding of a chloride ion in the lower part of the binding cavity likewise induces a Trp-out conformation, further underlining that the flexibility of this region is influenced by ligand or ion occupancy (PDB code: 1HKX)^6^. This prompted us to investigate the contribution of Trp403, first by mutating the CaMKIIα Trp into Leu (W403L), which is the residue present in the other CaMKII variants. This resulted in a slight reduction of [^3^H]HOCPCA binding in the W403L mutant (70% of WT, Fig. 2B) with unchanged overall protein expression (Fig. 2C). The reduction in binding indicates that Trp403 is a contributing conformational factor for binding of GHB ligands to CaMKIIα.

Despite the lack of Trp403 in CaMKIIβ crystal structures, alignment of beta sequences revealed a similar flexible loop (Fig. S1, Table S2). As such, we explored how introduction of a Trp at the corresponding location in CaMKIIβ (L467) might affect GHB ligand binding. To this end, [^3^H]HOCPCA binding was evaluated in CaMKIIβ via the single mutant L467W and the double mutant L467W/C533R. Neither of these led to any noticeable radioligand binding of [^3^H]HOCPCA (Fig. 2D) albeit protein expression was intact (Fig. 2E).

### Development of [^3^H]O-5-HDC as a tool to study compound-induced CaMKIIα conformations

Chemically, HOCPCA and 5-HDC are representatives of two different GHB ligand classes. Given their difference in size and binding kinetics (5-HDC has a lower off-rate that HOCPCA^14^), but also the fact that they stabilize different conformations of CaMKIIα; the Trp-in and Trp-out conformations, respectively^14^, makes them interesting to compare. Thus, based on the hypothesis that hub dynamics involving the Trp403 containing pocket loop could impact the selectivity of GHB ligands, we aimed to develop a radioligand analogue of 5-HDC that could serve as a useful tool to discern conformational effects. Since 5-HDC has been found to be chemically unstable^17^, the oxygen-bridged version (O-5-HDC) was used instead (Fig. 1A). These compounds were found to have similar K_i_ values in CaMKIIα binding assays previously reported (K_i_ of 5-HDC = 35 nM, O-5-HDC = 27 nM)^17^. As [^3^H]O-5-HDC is a novel tool, a synthesis overview is provided in Fig. 3A and full details in the Supporting Information. Notably, two tritium isotopes are incorporated per molecule of O-5-HDC, leading to a high specific activity of 48.2 Ci/mmol.

**Figure 3:**
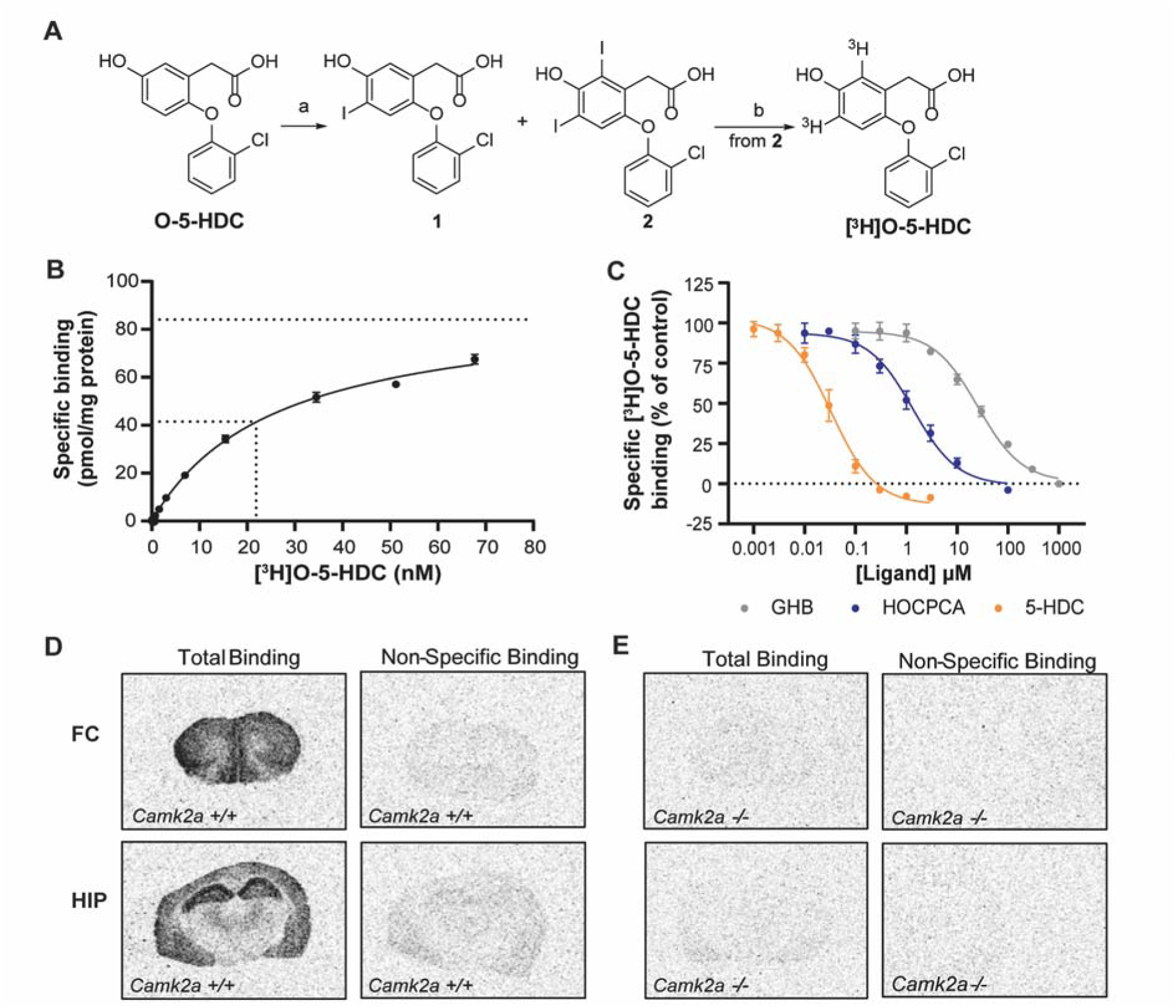
Development and characterization of [^3^H]O-5-HDC. **A)** Schematic overview of [^3^H]O-5-HDC synthesis. Reagents and conditions: a) I_2_, 30 % H_2_O_2_ (w/w%) aqueous solution, H_2_O 50 °C, overnight. (b) Pd/C, ^3^H_2_, Et_3_N, MeOH, 2 hours. **B)** Saturation isotherm of [^3^H]O-5-HDC; 1 mM GHB for NSB. **C)** Concentration-dependent inhibition of [^3^H]O-5-HDC binding by GHB analogues. **B,C)** Data are representative of at least three independent experiments performed in triplicates and shown as mean ± SD; K_i_ values are summarized in Table S4. Specific binding of [^3^H]O-5-HDC in frontal cortex (FC) and hippocampus (HIP) in brain slices from **D)** *Camk2a* +/+ mice and **E)** *Camk2a* -/- mice; 1 mM GHB for NSB. The autoradiograms are representative of four slices from a single experiment.

Before using [^3^H]O-5-HDC for addressing selectivity, assay conditions were established with native CaMKIIα in rat cortical homogenate (detailed in Supporting Information, Fig. S2 and Table S3). The developed assay is highly similar to the protocol for [^3^H]HOCPCA using rat cortical homogenate^22^. The binding reaction followed a saturable profile (Fig. 3B, Table S4) and revealed a K_D_ value of 22 nM and a B_max_ of 86.5 pmol/mg protein. This is noteworthy, as the K_D_ signifies a 10-fold improved affinity compared to [^3^H]HOCPCA (Table S4)^22,26^. Moreover, K_i_ values for GHB, HOCPCA and 5-HDC were determined, yielding values of 24.1 µM, 1.27 µM and 0.030 µM, respectively (Fig. 3C, Table S5). Interestingly, the smaller sized ligands HOCPCA and GHB showed 10-fold lower affinity in displacing [^3^H]O-5-HDC compared to [^3^H]HOCPCA (Table S5). Notably, with 1 mM GHB to determine NSB, 5-HDC displayed a plateau below 0% (around -10 %). On the contrary, inhibition of [^3^H]HOCPCA binding by 5-HDC revealed a bottom plateau around 20%^14^. This clearly highlights small but significant probe difference in two radioligands, despite interaction with the same core residues. As such, this may reflect that different, or more, molecular interactions are at play for the slightly larger compound O-5-HDC. Alternatively, it could reflect altered levels of occupancy, although B_max_ values are very similar for [^3^H]HOCPCA and [^3^H]O-5-HDC^14^. To confirm CaMKIIα selectivity for [^3^H]O-5-HDC as for [^3^H]HOCPCA, autoradiography on mouse brain slices using 0.09 nM [^3^H]O-5-HDC was conducted. This revealed high density binding in frontal cortex and hippocampus (Fig. 3D), matching the regional binding of [^3^H]HOCPCA^27^. The observed binding was displaceable by 1 mM GHB and was absent in *Camk2a^-/-^*(KO) mice (Fig. 3E), confirming specific CaMKIIα interaction. Compared to [^3^H]HOCPCA, the use of [^3^H]O-5-HDC resulted in autoradiograms of higher quality due to the higher radioligand sensitivity.

### Successful introduction of [^3^H]O-5-HDC binding into CaMKIIβ

With the availability of [^3^H]O-5-HDC as a conformationally selective and highly specific CaMKIIα radioligand, we revisited our mutants in the new binding assay. As expected, [^3^H]O-5-HDC binding to the mutants of the core molecular determinants R453Q/R469Q was still completely absent (P < 0.0001) (Fig. 4A). Furthermore, as for [^3^H]HOCPCA, we observed a slight, significant reduction in specific binding of [^3^H]O-5-HDC to W403L (75% of WT, P = 0.0179) (Fig. 4A). This coincides with maintained binding of another GHB analogue, PIPA, to purified CaMKIIα-W403L hub protein in surface plasmon resonance studies with similar, yet slightly different K_D_ values (1.4 µM for CaMKIIα WT and 2.8 µM for CaMKIIα-W403L)^19^.

**Figure 4:**
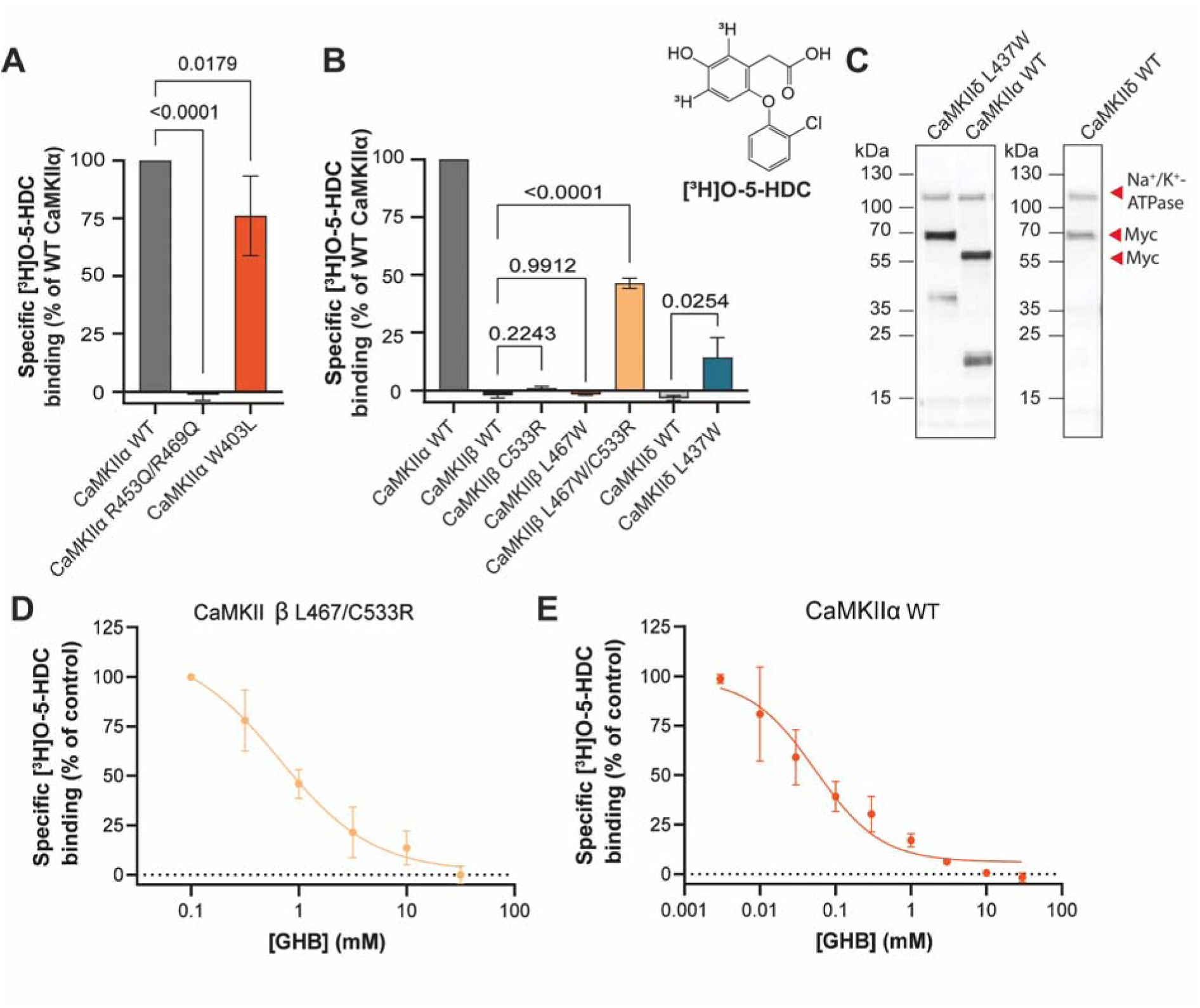
Mutation of residue 467 (L467W) in the CaMKIIβ pocket loop promotes binding. **A)** Specific binding of [^3^H]O-5-HDC to the CaMKIIα mutant R453Q/R469Q and W403L. **B)** Specific [^3^H]O-5-HDC binding to CaMKIIβ C533R, L467W and L467W/C533R and CaMKIIδ L437W mutants. **C)** Western blot validation of protein expression; Na^+^/K^+^-ATPase as loading control **D)** Concentration-dependent displacement of [^3^H]O-5-HDC binding by GHB in CaMKIIβ L467W/C533R, and **E)** CaMKIIα WT. Data are pooled from three to four independent experiments performed in technical triplicates and shown as mean ± SD. One Way ANOVA, post hoc Dunnett’s test; significance level P<0.05.

Finally, [^3^H]O-5-HDC binding was performed to the CaMKIIβ mutants, L467W, C533R and L467W/C533R. Whereas no binding was observed for the single mutations, the double mutant L467W/C533R now presented with a significant amount of specific [^3^H]O-5-HDC binding compared to WT (P<0.0001) (Fig. 4B). Further, the specific binding at CaMKIIβ L467W/C533R was displaceable by GHB in a concentration-dependent manner (IC_50_ value of 710 µM (3.17 + 0.07) (Fig. 4D). However, GHB displayed a 10-fold decrease in affinity when compared to CaMKIIα WT (IC_50_ value of 61.7 µM (4.28 ± 0.20)) (Fig. 4E), suggesting involvement of, yet unidentified, additional residues or features. Curiously, introduction of the corresponding Trp mutation in CaMKIIδ (L437W) resulted in detectable [^3^H]O-5-HDC specific binding (Fig. 4B) (P = 0.025), however to a lesser extent than CaMKIIβ L467W/C533R (14% and 46% of CaMKIIα WT specific binding, respectively). This could be explained by the longer C-tail of CaMKIIδ, which could affect pocket accessibility dependent on the position of this flexible region in the upper part of the cavity.

Altogether, with the advent of [^3^H]O-5-HDC and carefully engineered mutations at key positions, we were able to introduce GHB ligand binding into CaMKIIβ and δ. One reason why [^3^H]HOCPCA binding was not achieved with CaMKIIβ could reflect the chemical differences of the radioligands, i.e. only O-5-HDC contains the functional groups (aromatic rings) to enable a different conformation of the pocket loop possibly as a result of Trp403 flip. As such, [^3^H]O-5-HDC might stabilize a conformation favourable for GHB analogue binding and alleviate the possibility of binding to otherwise non-binding CaMKII isozymes. Indeed, the absence of Trp403 in the CaMKIIα pocket decreases protein thermal stability^19^ and the pocket loop was previously suggested to mediate an interaction between the kinase domain and the hub domain in a docked state, which highlights a potential, yet unclear, contribution to physiological CaMKIIα function^7^. Additionally, with the decreased affinity observed in the CaMKIIβ mutant compared to CaMKIIα, [^3^H]HOCPCA binding may simply not be strong enough to be detected due to its lower affinity and faster off-rate. Thus, the molecular determinants for GHB ligand selectivity are not only driven by the four core binding pocket residues, but also by overall hub dynamics, signified by interactions with flexible regions located in the upper part of the hub cavity. Finally, additional factors such as other parts of the flexible region or the C-tail may impact selectivity.

## Conclusion

In summary, site-directed mutagenesis studies revealed a common set of core binding residues for two different GHB radioligands, as well as probe-dependent molecular determinants in the flexible upper hub cavity. The novel radioligand, [^3^H]O-5-HDC permitted introduction of binding in the otherwise non-binding isozyme, CaMKIIβ, and to a smaller extent, CaMKIIδ. We conclude that larger-type GHB analogues such as O-5-HDC are more likely to form stabilising interactions than smaller-type ligands such as GHB and HOCPCA by compensating for the Trp403 movement. These findings corroborate the CaMKII *alpha* selectivity of chemically diverse GHB analogues. In contrast to standard non-selective CaMKII ligands (e.g. CN21, AS100105 or KN93)^28^, it highlights the potential of GHB analogues for targeted CaMKIIα drug discovery.

## Abbreviations

BSA: bovine serum albumin;
CaM: calmodulin;
CaMKII: Ca^2+^/Calmodulin-dependent protein kinase II;
GHB: γ-hydroxybutyric acid;
HOCPCA: 3-hydroxycyclopent-1-enecarboxylic acid;
5-HDC: 5-hydroxydiclofenac;
KO: knock out;
NSB: non-specific binding;
O-5-HDC: oxygen-bridged 5-HDC;
TBS-T: Tris-buffered saline with Tween-20;
WT: wild type

## Acknowledgements

This work was supported by the Independent Research Fund Denmark (Grant numbers 8020-00156B and 1026-00335B to P.W.), the Lundbeck Foundation (R277-2018-260 to P.W.), Fougner Hartmann’s Family Foundation, and supported by the Czech Academy of Sciences (RVO: 61388963). We thank Dr. Sara Solbak for fruitful discussions.

