## Supplementary material for "Allosteric factors in the calcium/calmodulin-responsive kinase II hub domain determine selectivity of GHB ligands for CaMKIIα": SI file

**List of material included:**

**Fig. S1*:*** Crystal structure alignment of four CaMKIIβ structures

**Fig. S2:** Characterization of [^3^H]O-5-HDC binding to rat cortical homogenates

**Table S1:** Overview of the CaMKIIα crystal structures with PDB codes used for the overlay in Fig. 2A

**Table S2:** Overview of the CaMKIIβ, -γ and -δ crystal structures with PDB codes used for the overlay in Fig. S1

**Table S3:** Association and dissociation rate constants for [^3^H]O-5-HDC

**Table S5:** Competitive inhibition of [^3^H]O-5-HDC binding by GHB, HOCPCA and 5-HDC

### **Supporting methods:** Synthesis of [^3^H]O-5-HDC; Cell culture and transfections; Preparation of whole-cell homogenates and protein determination; Western blot analysis; Radioligand binding assay with native cortical membrane homogenates: Data analysis; Computational modelling.

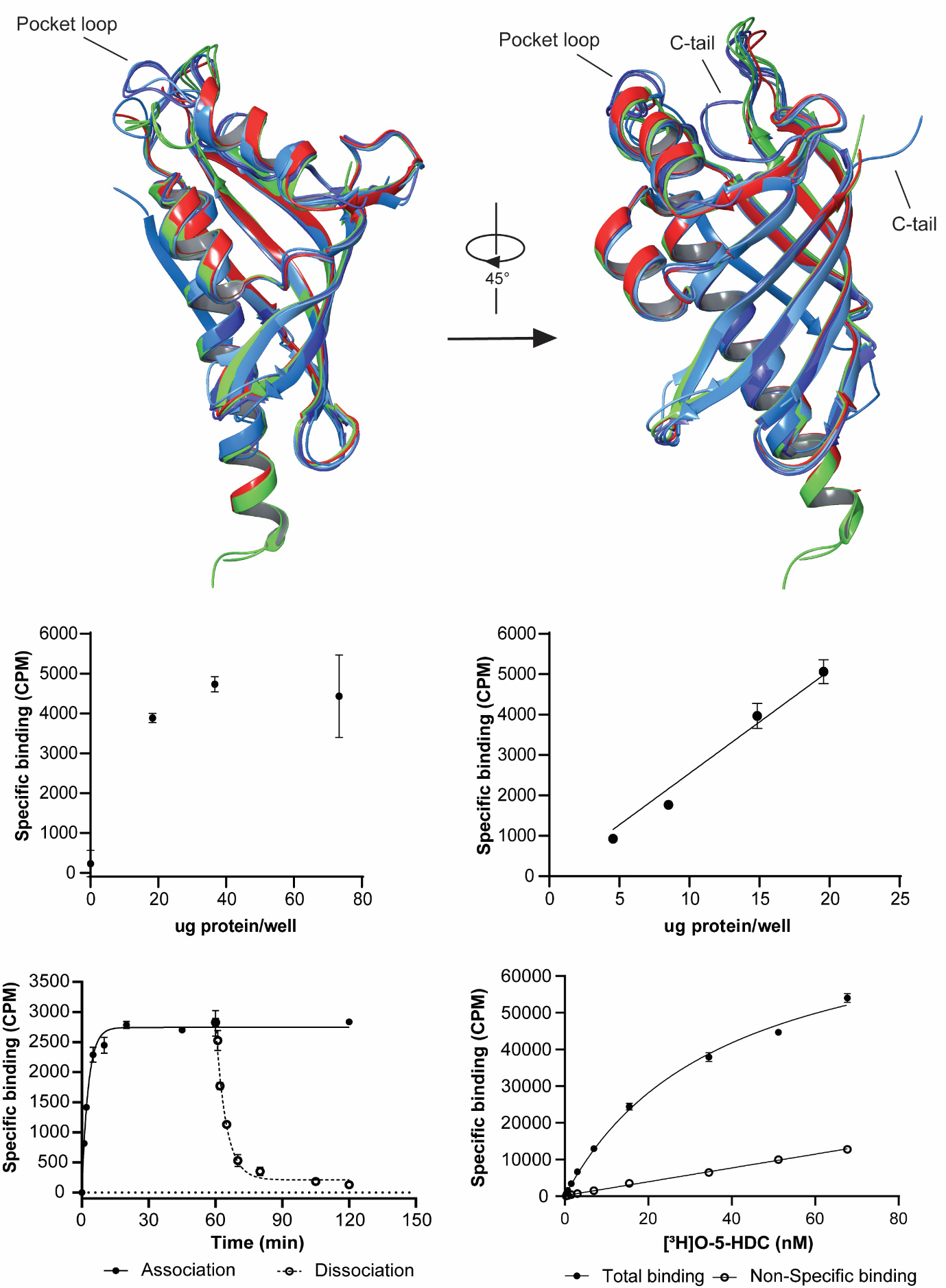

**Fig. S1:** **Crystal structure alignment of four CaMKIIβ structures** (PDB codes: 5IG5 in azure blue, 7URZ in faded azure blue, 7URY in faded blue and 7URW in blue), two CaMKIIγ structures (PDB codes: 2UX0 chain A and chain E in faded green) and one CaMKIIδ structure (PDB code: 2W2C red). The hub domain including the pocket loops and C-tails are depicted from two different angles. Note: two residues in the pocket loop of CaMKIIδ (PDB code: 2W2C) and between one to seven residues in the C-tail of multiple structures (PDB code: 5IG5, 7URZ, 2UX0 and 2W2C) were not resolved.

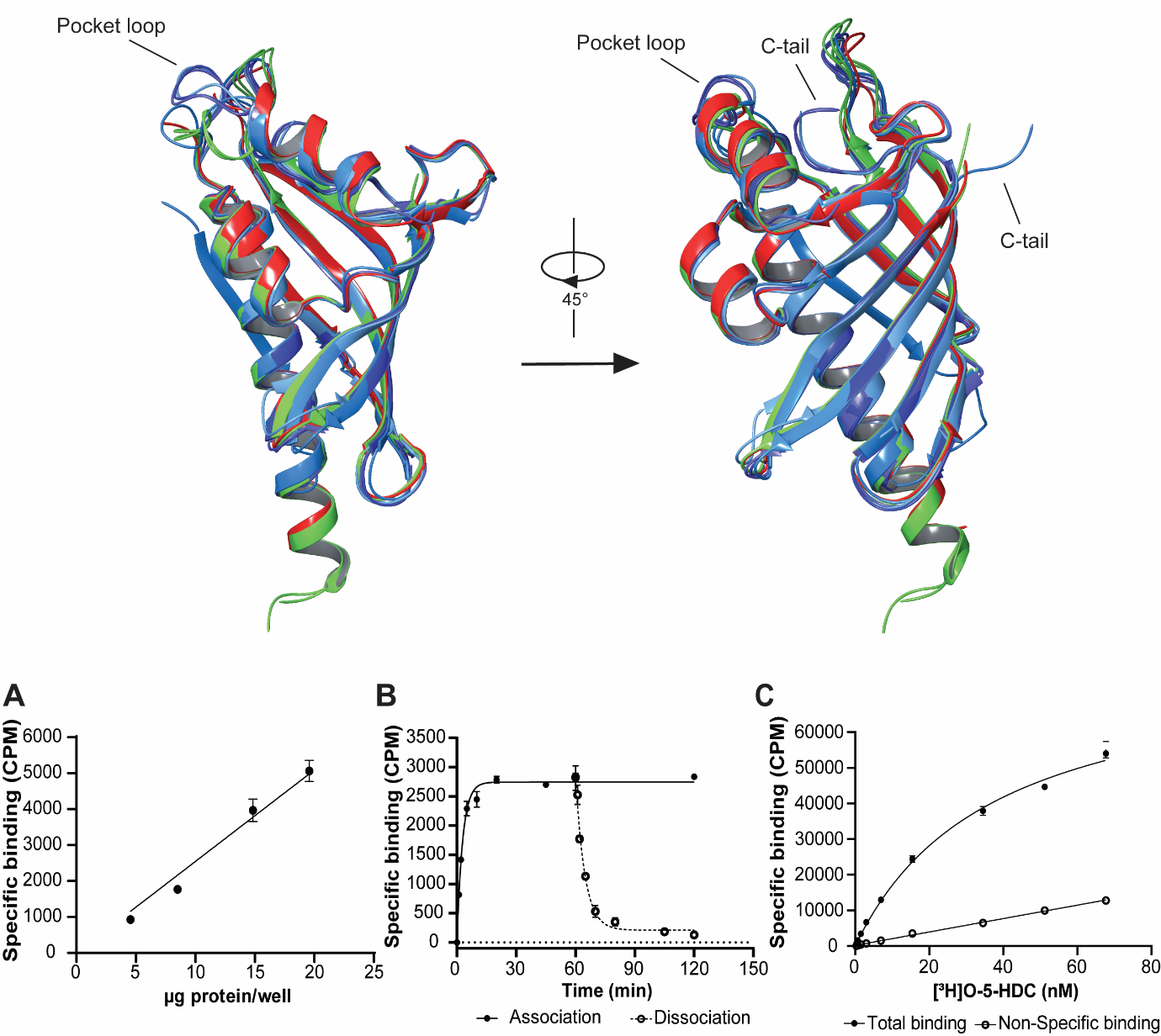

**Fig. S2: Characterization of [^3^H]O-5-HDC binding to rat cortical homogenates.**

**A)** Protein dependent increase of 3 nM radioligand binding with linearity up to approximately 20 µg protein/well (n = 2). **B)** Association and dissociation of 5 nM [^3^H]O-5-HDC. Dissociation was initiated after equilibrium (at 60 min) by addition of 1 mM GHB. (n = 2) **C)** Saturation binding profile of 0.2 to 75 nM of [^3^H]O-5-HDC. The non-specific binding obtained by 1 mM GHB displays a linear relationship. (n = 3) Data are representative of two to three independent experiments performed in technical triplicates and presented as mean ± SD**.**

**Table S1:** **Overview of the CaMKIIα crystal structures with PDB codes used for the overlay in Fig. 2A**

| PDB code | Chain | Unbound vs. bound (compound) | Colour |
| --- | --- | --- | --- |
| 7REC | A | Unbound | Blue |
| 7REC | B | Bound (5-HDC) | Red |
| 3SOA | A | Unbound | Dark blue |
| 6OF8 | D | Unbound | Light blue |
| 1HKX | A | Bound (Cl^-^) | Orange |

**Table S2:** **Overview of the CaMKIIβ, -γ and -δ crystal structures with PDB codes used for the overlay in Fig. S1**

| Isozyme | PDB code | Chain | Colour |
| --- | --- | --- | --- |
| CaMKIIβ | 5IG5 | F | Azure |
| CaMKIIβ | 7URZ | B | Faded azure |
| CaMKIIβ | 7URY | E | Faded blue |
| CaMKIIβ | 7URW | E | Blue |
| CaMKIIγ | 2UX0 | A | Faded green |
| CaMKIIγ | 2UX0 | E | Faded green |
| CaMKIIδ | 2W2C | H | Red |

**Table S3: Association and dissociation rate constants for [^3^H]O-5-HDC.** The on and off rates were estimated by fitting data to exponential association and dissociation. Data are stated as mean ± S.E.M. (n=2)

| Association | | Dissociation | |  |
| --- | --- | --- | --- | --- |
| K_obs_ (min^-1^) | **Half-time** (min) | **K_of_**_f_ (min^-1^) | **Half-life** (min) | **K_on_** (min^-1^M^-1^) |
| 0.43 ± 0.08 | 1.69 ±0.32 | 0.21 ± 0.00 | 3.31 ± 0.03 | 1.33*10^8^ ± 3.98*10^7^ |

**Table S4: Saturation data on [^3^H]O-5-HDC binding to rat cortical homogenates.** Data are based on three independent experiments performed in technical triplicates (n=3) with comparative values from [^3^H]HOCPCA binding^1^.

|  | [^3^H]O-5-HDC | Comparative values [^3^H]HOCPCA binding^1^ |
| --- | --- | --- |
| K_d_ (nM) (pK_d_ ± SEM) | 21.96 (7.7 ± 0.04) | 260 (6.6 ± 0.06) |
| B_max_ (pmol/mg protein ± SEM) | 86.5 ± 9.06 | 43 |

**Table S5: Competitive inhibition of [^3^H]O-5-HDC binding by GHB, HOCPCA and 5-HDC.** K_i_ values were derived using a one-site fit model and the Cheng-Prusoff equation (K_d_ = 21.96 nM). Data are based on four independent experiments performed in technical triplicates (n = 4) with comparative values from [^3^H]HOCPCA binding^1^.

| Compound | K_i_ (µM) (pKi ± S.E.M.) | Comparative K_i_ (µM) values [^3^H]HOCPCA binding^1^ |
| --- | --- | --- |
| GHB | 24.14 (4.63 ± 0.06) | 3.0 |
| HOCPCA | 1.27 (5.93 ± 0.12) | 0.13 |
| 5-HDC | 0.030 (7.55 ± 0.06) | 0.022 |

### **Supporting materials and methods**

**Synthesis of [^3^H]O-5-HDC, general procedure:** All reagents and solvents (reagent or chromatography grade) were purchased from commercial suppliers and were used without further purification unless specified. ^1^H or ^13^C-NMR spectra were recorded on a Bruker Advance 400MHz spectrometer assembled with a 5 mm BBFO probe or a Bruker Advance 600 MHz spectrometer with a cryogenically cooled 5 mm 13C/1H DCH probe at 300 K. ^3^H NMR spectra were recorded at 320 MHz, with a Bruker Avance II 300 MHz instrument at 298K. Data were tabulated in the following order: chemical shift (δ) [multiplicity (b, broad; s, singlet; d, doublet; t, triplet; q, quartet; m, multiplet), coupling constant(s) J (Hz), number of protons]. Methanol-d_4_ (reference signals δ_H_ = 3.31 ppm, δ_C_ = 49.00 ppm) was used as the solvent for NMR analysis, and the solvent residue peak was used as the internal reference.

Purification of non-radioactive compounds was carried out using preparative reverse-phase HPLC on an UltiMate HPLC system (Thermo Scientific) consisting of an HPG-3200BX pump, a Rheodyne 9725i injector, a 10 mL loop, an MWD-3000SD detector (254nm), and an AFC-3000SD automated fraction collector using a Gemini-NX C18 column (21.2 × 250 mm, 5 μm, 110 Å) (Phenomenex). For HPLC control, data collection, and data handling, Chromeleon software ver. 6.80 was used. UPLC-MS analysis was carried out on a Waters Acquity H-class UPLC with a Sample Manager FTN and a TUV dual wavelength detector coupled to a QDa single quadrupole analyzer using electrospray ionization (ESI). UPLC separation was achieved with a C18 reversed-phase column (Acquity UPLC BEH C18, 2.1 mm × 50 mm, 1.7 μm) operated at 40 °C. Data acquisition was controlled by MassLynx ver. 4.1 and data analysis was done using Waters OpenLynx browser ver. 4.1.

HPLC separation of [^3^H]O-5-HDC was performed on the Alliance e2695 module equipped with 2998 PDA detector (Waters, Milford, MA, USA) and operated by the Empower 3 Pro software. Radiochromatograms were recorded by the β–radioactivity HPLC flow detector Ramona Star equipped with the LS-pump (Elysia-Raytest, Straubenhardt, Germany). Liquid scintillation measurements were done on the Tri-Carb 2900 liquid scintillation counter (Perkin Elmer, Downers Grove, IL, USA) in a Rotiszint® eco plus cocktail. Mass spectra were obtained by LC-MS analysis. LC-MS was performed using an Agilent 1260 HPLC system with a Waters Acquity UPLC BEH 1.9 μm C-18 column (100 × 2.10 mm) using a linear gradient elution from buffer A (H_2_O:TFA, 100:0.1 v/v%) to buffer B (MeCN:TFA, 100:0.1 v/v%) from 2% to 100% B over 6 min, maintaining a flow rate of 0.3 mL⁄min, coupled to an Agilent 6230 TOF series mass spectrometer with an electrospray ionization source.

**Synthesis of [^3^H]O-5-HDC:** The [^3^H]O-5-HDC radioligand was synthesized from its non-radioactive counterpart, O-5-HDC^2^, using a two-step synthetic strategy (Fig. 3A). Facilitated by the phenolic group, aromatic iodination of O-5-HDC using elemental iodine produced the di-iodinated precursor **1**. Compound **1** was subsequently subjected to a palladium-catalyzed tritiodehalogenation reaction under atmospheric tritium gas (8.5 Ci, 310 GBq) for 2 hours, permitting the introduction of tritium isotope to the core O-5-HDC scaffold^3^. To this end, a total amount of 31 mCi of [^3^H]O-5-HDC was prepared, exhibiting a high specific activity of 48.2 Ci/mmol (1.8 TBq/mmol) and a radio-chemical purity of 99%. Finally, [^3^H]O-5-HDC was formulated to 1 mCi/mL in ethanol (UV grade) for further pharmacological studies.

Importantly, the tritiodehalogenation approach demonstrated a privileged reactivity of aryl iodide over aryl chloride present in compound **1** during competitive reductive dehalogenation and a high regioselectivity^3,4^. This is evidenced by the absence of dechlorinated by-products at the given reaction time and the exclusive incorporation of tritium at the desired labeling positions in [^3^H]O-5-HDC.

In comparison to O-5-HDC, the iodoarenes **1** and **2** displayed a 100X increase in affinity to mid-micromolar concentrations when evaluated in [^3^H]NCS-382 binding to rat brain cortical homogenates as previously reported^2,5^. This reveals an extremely tight fit and limited space around such iodinating positions in the binding pocket, where introduction of iodine to the *ortho*-positions of the phenol ring in O-5-HDC proved detrimental to the affinity.

**Synthesis of iodoarene (1).** To a suspension of O-5-HDC (0.11 g, 0.38 mmol^2^) and I_2_ (0.15 g, 0.58 mmol) in distilled water (2 mL) was added 30% H_2_O_2_ aqueous solution (0.12 mL, 1.2 mmol). The reaction mixture was stirred overnight at 50ºC in the dark and quenched by saturated Na_2_S_2_O_3_ solution. The aqueous phase was extracted with EtOAc three times. The combined organic phases were dried over Na_2_SO_4_, filtered, and evaporated in vacuo. Purification by preparative HPLC (gradient 30−60% B, eluent A (H_2_O/TFA, 100:0.1) and eluent B (MeCN/H_2_O/TFA, 90:10:0.1) at a flow rate of 20 mL min^−1^, over 15 min) furnished **1** (47 mg, 23%) as a white solid. ^1^H NMR (400 MHz, Methanol-d_4_) δ 7.49 (dd, J = 7.9, 1.6 Hz, 1H), 7.27 (td, J = 7.8, 1.7 Hz, 1H), 7.13 (td, J = 7.7, 1.5 Hz, 1H), 7.10 (s, 1H), 6.93 (dd, J = 8.1, 1.5 Hz, 1H), 3.95 (s, 2H). ^13^C NMR (101 MHz, Methanol-d_4_) δ 173.4, 153.9, 153.7, 149.8, 132.7, 131.9, 129.5, 128.9, 126.1, 126.0, 121.0, 95.5, 83.5, 42.0. UPLC-MS: m/z calculated [M-H]^-^ for C_14_H_8_ClI_2_O_4_ = 528.82, found [M-H]^-^ = 528.8 and [M-H+2]^-^ = 530.9.

**Preparation of [^3^H]O-5-HDC.** Compound **1** (5.5 mg, 10.4 μmol), Pd/C (11 mg, 10.4 μmol, 10% w/w% loading), Et_3_N (15.0 μL, 106.0 μmol) and MeOH (0.5 mL) were added to a round bottom flask (1 mL). The tritiodeiodination was carried out on the customized stainless tritium manifold system (RC Tritec, Teufen, Switzerland) in the glove box. The mixture was degassed by successively freezing-thawing (liquid N_2_ and vacuum of turbopump) three times. Career-free ^3^H_2_ gas (15.9 Ci, 590 GBq) stored on a uranium bed as uranium tritide was released (966 mbar) by heating to 500 °C, and then was directed into the reaction mixture. ^3^H_2_ gas (8.5 Ci, 310 GBq) was absorbed on the Pd/C catalyst. The reaction mixture was stirred vigorously at room temperature for 2 hours, and then frozen down using liquid N_2_. The unreacted ^3^H_2_ gas was back-trapped by the waste tritium uranium bed. The mixture was filtrated through a 0.45 μm PTFE syringe filter, and the reaction flask and filter were washed with 1mL MeOH three times. The labile activity was removed by repeatedly lyophilizing the added MeOH (3×5 mL) into the mixture. The obtained crude was re-taken into a mixed solution of 4 mL MeCN/H_2_O (60:40) and 1 mL MeOH. One-fifth of the solution was purified using preparative HPLC, giving 31 mCi of [^3^H]O-5-HDC with a specific activity of 48.2 Ci/mmol (1.8 TBq/mmol). Finally, **[**^3^H]O-5-HDC was formulated into 1.0 mCi/mL in EtOH (UV grade). ^3^H (^1^H decoupled) NMR (320 MHz, Methanol-d_4_) δ 6.89 – 6.88 (m, 1[^3^H]), 6.72– 6.71 (m, 1[^3^H]). ^3^H (^1^H coupled) NMR (320 MHz, Methanol-d_4_) δ 6.88 (d, J = 3.1 Hz, 1[^3^H]), 6.71 (dd, J = 9.2, 3.1 Hz, 1[^3^H]). LC-MS: m/z calculated [M-H]^-^ for C_14_H_8_^3^H_2_ClO_4_ = 281.0, found [M-H]^-^ = 281.0 and [M-H+2]^-^ = 283.0. Radiochemical purity (R.C.P.) by anal. radio-HPLC: 98.7% (254 nm).

### **Cell culture and transfections:** HEK293T cells were purchased from ATCC (293T-ATCC; #CRL-3216; authenticated to be mycoplasma-free) and maintained in Dulbecco’s Modified Eagle Medium (DMEM) with GlutaMax medium (#61965026, Gibco, Thermo Fisher Scientific, West Palm Beach, FL, USA), supplemented with 10% (v/v) fetal bovine serum (Gibco) and 1% (v/v) penicillin/streptomycin (#15140122, Invitrogen). Cells were kept at 37 °C in a humidified atmosphere and 5% CO_2_. For transfection, we used polyethyleneimine (PEI, #23966 from Polysciences Inc., Warrington, PA, USA) in a DNA:PEI ratio of 1:3 similar to^6^. On the day before transfection, 4.5x10^6^ cells were plated out in 15 cm cell culture dishes. For transfection, 16 µg DNA was mixed with 48 µg PEI and diluted in 2 mL DMEM + 1% (v/v) penicillin/streptomycin. After 15 min incubation at room temperature, the DNA/PEI mixture was added to the cells.

### **Preparation of whole-cell homogenates and protein determination:** Cells were harvested 48h post-transfection using ice-cold DPBS (Gibco) and cell scrapers. Cells were centrifuged at 1,500 x *g* for 10 min, and the resulting cell pellet was re-suspended in ice-cold binding buffer (50 mM KH_2_PO_4_, pH 6.0). The cells were homogenised using 2 x 1 mm zirconium beads in a Bullet Blender (NextAdvance, NY, USA) for 20 sec at maximum speed. Protein concentration was determined using Bradford Protein assay (Bio-Rad Laboratories, Copenhagen, Denmark) according to manufacturer’s instructions.

### **Western blot analysis:** Based on the protein determination, equal amounts of protein from whole-cell homogenates were mixed with 4x NuPage® LDS Sample Buffer 4x (ThermoFisher Scientific) and 100 mM DL-dithiothreitol (Sigma-Aldrich) followed by heating for 10 min at 40 °C, sonication and centrifugation for 2 min at 4 °C at 11,000 g. For the western blot analysis, 5 µg protein was loaded into 4–20% Mini-PROTEAN® TGX™ Precast Protein Gels (Bio-Rad) followed by transfer onto polyvinylidene difluoride membrane (Bio-Rad) using semi-dry TransBlot® Turbo^TM^ Transfer System (Bio-Rad) according to the manufacturer’s instructions. Membranes were blocked for 1 h at room temperature in 3% bovine serum albumin (BSA) dissolved in Tris-Buffered Saline + 0.05 % Tween-20 (TBS-T). The primary antibodies targeting either c-myc (Mouse monoclonal, Invitrogen, 1:1000 dilution) or native CaMKIIβ (mouse monoclonal, Invitrogen, 1:2000 dilution) co-incubated with the antibody for Na^+^/K^+^-ATPase as loading control (rabbit monoclonal, Abcam, 1:10,000 dilution) for 1 hour at room temperature was followed by 3 x 5 min wash in TBS-T. Likewise, the species-specific secondary antibodies conjugated with horseradish peroxidase (goat polyclonal anti-mouse and -rabbit, Agilent, 1:2000) co-incubated for 1 hour at room temperature followed by 3x 10 min wash in TBS-T. All antibody dilutions were prepared using 1 % BSA in TBS-T, stored at -20 °C (primary) or 4 °C (secondary) and re-used up to four times. Membranes were probed for up to 4 min with 1:1 mixture of the substrate, ECL Detection Reagent (GE Healthcare). The luminescence signals were captured using either iBright^TM^ FL1500 Imaging System (Invitrogen) or FluorChem HD2 (Alpha Innotech).

**Radioligand binding assay with native cortical membrane homogenates:** For equilibrium binding, incubation of 10-15 µg protein on ice for 60 min with 5 nM [^3^H]O-5-HDC in a 50 mM KH_2_PO_4_, pH 6.0 buffer (total volume of 200 µl) was used. Binding was found to be linear with 0-25 ug total protein/well. Kinetics was assessed with association and dissociation curves using time points 1-120 min, and [^3^H]O-5-HDC saturation using concentrations of 0.5-100 nM. In all experiments, NSB was determined with 1 mM GHB. Protein-ligand bound complexes were collected by rapid filtration through GF/C filter plates and counts per minute (CPM) measured in a TopCount NXT Microplate Scintillation counter (PerkinElmer) with 3 min counting per well. For saturation experiments, CPM was converted to DPM via a standard quench curve. Check of exact radioligand concentrations were determined with the TriCarb and no ligand depletion was observed.

**Data analysis:** The specific binding was determined as Total binding substracted NSB. For the pooling of data from recombinant binding, the specific binding of the variant was normalised to the specific binding of CaMKIIα WT within each experiment. Binding displacement curves were analysed using One Site-Fit logIC50 model, and the IC_50_ values of GHB determined using the following equation:

$$Specific binding (\% of control)=\frac{Top-Bottom}{1+{10}^{(\log(IC_{50}-[compound]))}}$$

where *top* and *bottom* represent the upper and lower plateaus of the curve, respectively. *IC_50_* is the concentration of the inhibitor that inhibits 50 % of the response interval between minimum and maximum response. The *[compound]* is the logarithmic value of the applied GHB concentration.

K_i_ values were obtained from the One Site-Fit K_i_ model utilizing the following Cheng-Prusoff equation:

$$K_{I}=\frac{IC50}{1+{\left[ \mathrm{RL} \right]/K_{D}}}$$

Where [RL] represents the specific radioligand concentration from each experiment and KD is the dissociation constant of [^3^H]O-5-HDC acquired from saturation curves.

In saturation experiments, specific binding was corrected for NSB of each radioligand concentration and the resulting data were fitted to a one-site model by non-linear regression:

$$Specific bound ligand=\frac{B_{max}}{1+K_{D}/[L]}$$

Where K_D_ is the dissociation constant and B_max_ is the maximum specific binding.

**Computational modeling:** Alignment of CaMKIIβ, -γ and -δ was performed using the Maestro Schrödinger package version 13.2 (Schrödinger Release 2022-2: Maestro, Schrödinger, LLC, New York, NY, 2021). The crystal structures were prepared using the Protein Preparation Wizard^7^ (Schrödinger Release 2022-3: Protein Preparation Wizard; Epik, Schrödinger, LLC, New York, NY, 2021; Impact, Schrödinger, LLC, New York, NY; Prime, Schrödinger, LLC, New York, NY, 2021) including hydrogen optimisation by PROPKA at pH 7.4 and energy minimization using OPLS4 forcefield. The protein alignments of the listed structures were performed using the Protein Superimposition module by aligning the backbone atoms of PDB entry 5IG5 (largest RMSD = 1.488 Å).
